# Impact of Peripheral Auditory Structure on the Development of Auditory-Language Network in Children with Profound Sensorineural Hearing Loss

**DOI:** 10.1101/2023.02.09.527841

**Authors:** Yaoxuan Wang, Mengda Jiang, Yuting Zhu, Lu Xue, Wenying Shu, Xiang Li, Hongsai Chen, Yinghua Chu, Yang Song, Xiaofeng Tao, Zhaoyan Wang, Hao Wu

## Abstract

Profound congenital sensorineural hearing loss (SNHL) prevents children from developing spoken language. Cochlear implantation and auditory brainstem implantation can provide hearing sensation, but language development outcomes can vary, particularly for patients with inner ear malformations and/or cochlear nerve deficiency (IEM&CND). Currently, the peripheral auditory structure is evaluated through visual inspection of clinical imaging, but this method is insufficient for surgical planning and prognosis. The central auditory pathway is also challenging to examine in vivo due to its delicate subcortical structures. Previous attempts to locate subcortical auditory nuclei using fMRI responses to sounds are not applicable to deaf patients. In this study, we developed a new pipeline for mapping the auditory pathway using structural and diffusional MRI. We used a fixel-based approach to investigate the structural development of the auditory-language network for profound SNHL children with normal peripheral structure and those with IEM&CND under six years old. Our findings indicate that the language pathway is more sensitive to peripheral auditory condition than the central auditory pathway, highlighting the importance of early intervention for profound SNHL children to provide timely speech inputs. We also propose a comprehensive pre-surgical evaluation extending from the cochlea to the auditory-language network, which has promising clinical potential.

## Introduction

Congenital profound hearing loss (hearing level >90 dB diagnosed at birth or in early childhood) deprives children of spoken language development and has lifelong consequences for education, employment, and psychosocial status (Kral & O’Donoghue, 2010). The prevalence of neonatal hearing loss has increased from 1.09 to 1.7 cases per 1000 live births in the past two decades (*2019 CDC Early Hearing Detection and Intervention (EHDI) Hearing Screening & Follow-up Survey (HSFS)*. 2021; Alicia & Marcus, 2010). Most congenital hearing loss is caused by sensorineural impairments in the inner ear, cochlear nerve, and/or central auditory pathway (Korver et al., 2017). Approximately 15% to 39% of children with sensorineural hearing loss (SNHL) have bony inner ear malformations and/or cochlear nerve deficiency that can be identified with CT and MRI (Li et al., 2011; Mafong et al., 2002); the remaining 60% to 80% of cases are caused by membranous inner ear malformations, which are cellular-level abnormalities (Sennaroglu & Saatci, 2002).

Cochlear implantation (CI) and auditory brainstem implantation (ABI) are currently the only solutions for profound SNHL. CI directly stimulates the spiral ganglion cells, the first-order neurons of the auditory pathway, while ABI bypasses the cochlear nerve and stimulates the second-order auditory neurons in the cochlear nucleus when complex inner ear malformation or cochlear nerve aplasia makes CI inapplicable (Chen & Oghalai, 2016). Both CI and ABI are capable of providing hearing sensation and language development for prelingually deaf children; ABI recipients generally have poorer speech recognition performance and delayed and incomplete language development compared to CI recipients (Sennaroğlu et al., 2016). However, postoperative outcomes vary among individuals, and sometimes patients face the dilemma of choosing between CI and ABI when the prognosis of CI is not necessarily better than that of ABI, such as in cases of cochlear nerve hypoplasia (the cochlear nerve is present but thin) (Freeman & Sennaroglu, 2018). Thus, contemporary clinical imaging is insufficient for evaluating prognosis and assisting surgical strategies.

Speech is transmitted and processed through the auditory pathway and understood at the language network (Friederici, 2011). For prelingually deaf children, the deprivation of auditory inputs into this workflow may affect the structural and functional development of associated brain regions, the degrees of impairment of which may be relevant to later auditory and language performance after surgical reconstruction of hearing. Diffusion tensor imaging (DTI) studies have shown that children with profound SNHL exhibit decreased fractional anisotropy (FA) values along almost the entire auditory pathway, particularly in the inferior colliculus, medial geniculate body, and auditory radiation (Tarabichi et al., 2018). Additionally, hearing and language outcomes (categories of auditory performance) six to twelve months following CI were positively correlated with preoperative auditory pathway FA values (Chang et al., 2012; Huang et al., 2015; H. Wang et al., 2019; Wu et al., 2016). Similarly, brain structures associated with the language pathway, including the superior temporal gyrus (STG), Broca’s area, superior longitudinal fasciculus (SLF), and uncinate fasciculus (UF), also showed decreased FA values in children with profound hearing loss (H. Wang et al., 2019; S. Wang et al., 2019; Wu et al., 2016). The FA value of Broca’s area was positively correlated with speech recognition performance after CI implantation (Chang et al., 2012). These results indicate that children with profound SNHL have a delayed or impaired microstructural organization in the central auditory pathway and the language pathway that may affect subsequent implantation outcomes.

However, there are several challenges in this field. Firstly, the human auditory pathway is difficult to inspect non-invasively due to its delicate nodes connected by curving and crossing fibres, especially the parts that are buried deep in the brainstem (Zanin et al., 2019). Functional MRI (fMRI) responses to natural sounds have been used to locate subcortical auditory structures (Sitek et al., 2019), but this method is not applicable to patients with profound hearing loss. In fact, few studies have located the auditory brainstem of children with profound hearing loss, particularly the cochlear nucleus that ABI targets. Therefore, new localization and tractography methods are needed to precisely map the auditory pathway in vivo for individuals with normal or impaired hearing. Secondly, although earlier studies reported several altered fibre tracts related to language function, there has been insufficient investigation into the language pathway. Its component streams, which are segmented on both structural and functional bases, do not correspond to the major fibre tracts typically studied in whole-brain analyses. Finally, the diffusion tensor model used in these studies performs poorly in estimating regions with crossing fibres (Farquharson et al., 2013), which may lead to problematic tractography (particularly in the auditory pathway) and false interpretations of FA alterations in children with profound hearing loss. To overcome this methodological limitation, the present study implemented a state-of-the-art fixel-based analysis (FBA) approach (Dhollander et al., 2021). A “fixel” refers to an individual fibre population within a voxel, allowing for the quantification of white matter properties in fibre crossing areas.

Furthermore, earlier neuroimaging studies of profound congenital SNHL excluded children with bony inner ear malformations and/or cochlear nerve deficiency. However, these subjects comprise a significant proportion of patients with congenital hearing loss, and are more inclined to be faced with difficult surgical decisions and unsatisfactory post-implantation outcomes. Therefore, it is important to include and focus on children with auditory system malformations when studying central adaptations associated with profound hearing loss.

In the present study, we introduced a new pipeline for reconstructing the human auditory pathway, and examined the brain structural development of children with profound congenital sensorineural hearing loss at both the acoustic processing level and the speech perception level. Specifically, we included deaf children under six years old, with and without inner ear malformations and cochlear nerve deficiency, as well as normal hearing controls. We segmented the subcortical auditory nuclei using super-resolution track density imaging (TDI) maps and T1-weighted images, and tracked the auditory pathway and the language pathway using probabilistic tractography. Then, we used fixel-based analysis (FBA) to investigate the fibre properties of these two pathways. We also aimed to investigate whether the presence of inner ear malformations and/or cochlear nerve deficiency in patients would affect the structural development of their auditory-language networks. By doing so, we hope to provide new insights into the surgical strategies and rehabilitation of children with profound congenital SNHL.

## Results

### Demographics

Twenty-three children aged under six years old including thirteen patients with bilateral profound congenital sensorineural hearing loss (mean [SD], age, 30.92 [6.115] months; 9 males, 4 females) and ten normal hearing volunteers (mean [SD], age, 42.90 [4.270] months; 5 males, 5 females) matched on age and sex were included (Mann-Whitney U test: p-value for age = 0.077, p-value for sex = 0.446). Nearly half of the patients (six out of thirteen) had inner ear malformations and cochlear nerve deficiencies: two of them underwent ABI surgeries; the other four were ABI candidates. The remaining half exhibited normal structures and underwent CI surgeries (see ***Table 1***).

**Table 1.** Demographic information for patients with congenital bilateral profound sensorineural hearing loss.

| Sex | Age (months) | Gestational weeks | Birth weight (kg) | Inner ear structure (CT) | Cochlear nerve (CISS) | Surgery |
| --- | --- | --- | --- | --- | --- | --- |
| M | 26 | 39 | 3.25 | Cochlear aplasia | Deficiency | ABI |
| M | 24 | 39 | 2.7 | Normal | Normal | CI |
| F | 56 | 39 | 3.2 | Normal | Normal | CI |
| F | 65 | 39 | 2.9 | Normal | Normal | CI |
| M | 17 | 40 | 3.4 | Cochlear hypoplasia | Deficiency | ABI candidate |
| M | 32 | 40 | 3.3 | Normal | Normal | CI |
| M | 29 | 40 | 3 | Cochlear hypoplasia (L), Cochlear aplasia (R) | Deficiency | ABI |
| M | 14 | 40 | 2.9 | Normal | Normal | CI |
| M | 6 | 39 | 4.3 | Hypoplastic cochlear aperture | Deficiency | ABI candidate |
| F | 72 | 36 | 2.2 | Hypoplastic cochlear aperture | Deficiency | ABI candidate |
| M | 44 | 40 | 3.97 | Normal | Normal | CI |
| F | 9 | 40 | 3.6 | Normal | Normal | CI |
| M | 8 | 38 | 6 | Hypoplastic cochlear aperture | Deficiency | ABI candidate |

Inner ear structure and cochlear nerve were inspected using temporal bone High-Resolution CT and CISS MRI. The presence of inner ear malformation and cochlear nerve deficiency were determined based on Sennaroglu classification criteria (Sennaroglu et al., 2017); descriptions refer to a bilateral condition if the side (such as L or R) is not specified.

### Segmentation of subcortical auditory regions and tractography of the central auditory pathway

All subcortical auditory nuclei, including the bilateral cochlear nucleus (CN), superior olivary complex (SOC), inferior colliculus (IC), and medial geniculate body (MGB), were segmented in both group-average space and individual space with good inter-rater reliability (see ***Figure 1***).

**Figure 1.**
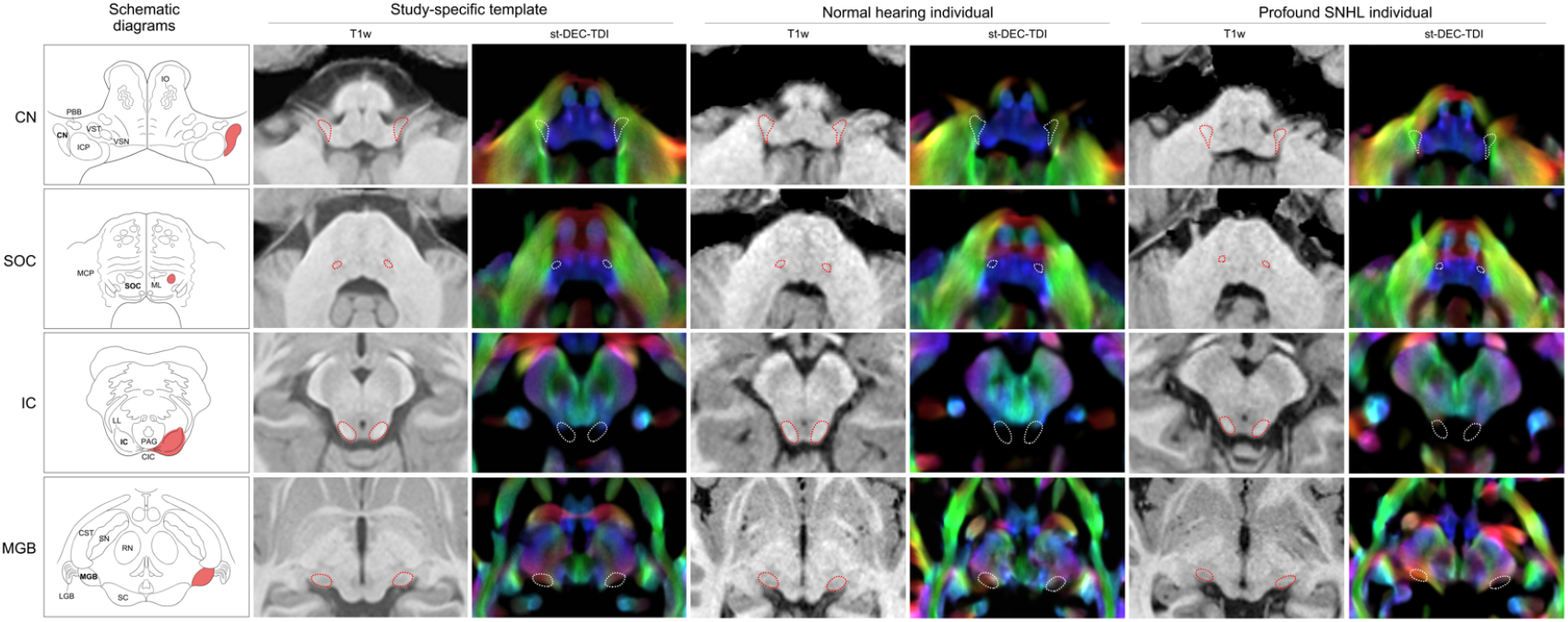
Segmentation of the subcortical auditory regions at group and individual level. Subcortical auditory regions are exhibited as red areas in the schematic diagrams and dotted lines in T1w images and st-DEC-TDI maps. All images are in the axial plane. Schematic diagrams for the CN, SOC, and IC were redrawn from *Moore (1987)* and for the MGB from *Duvernoys Atlas of the Human Brain Stem and Cerebellum (2009)*. Group-level segmentation was performed in the study-specific template averaged from all ten normal-hearing controls. Images from one normal hearing control (male, 32 months old) and one profound SNHL patient (male, 32 months old) were selected to show individual-level segmentation. Abbreviations: CN, cochlear nucleus; ICP, inferior cerebellar peduncle; PBB, ponto bulbar body; VSN, trigeminal spinal nucleus; VST, trigeminal spinal tract; IO, inferior olive; SOC, superior olivary complex; ML, medial lemniscus; MCP, medial cerebellar peduncle; IC, inferior colliculus; CIC, commissure of IC; LL, lateral lemniscus; PAG, periaqueductal grey; MGB, medial geniculate body; LGB, lateral geniculate body; SC, superior colliculus; RN, red nucleus; SN, substantia nigra; CST, corticospinal tract; St-DEC-TDI, short-tracks directionally encoded colour track density imaging.

We reconstructed the auditory pathway in vivo using probabilistic tractography in four subdivisions: the trapezoid body (TB), lateral lemniscus (LL), brachium of inferior colliculus (BIC), and acoustic radiation (AR) (see ***Figure 2***). The auditory pathway was generally symmetric bilaterally. Except for the fibres connecting the CN to the contralateral SOC, contralateral probabilistic tracking resulted in fewer streamlines than ipsilateral counterparts (fibres tracking from the CN or SOC to the contralateral IC showed fewer than 30 streamlines each and were thus removed from further analyses).

**Figure 2.**
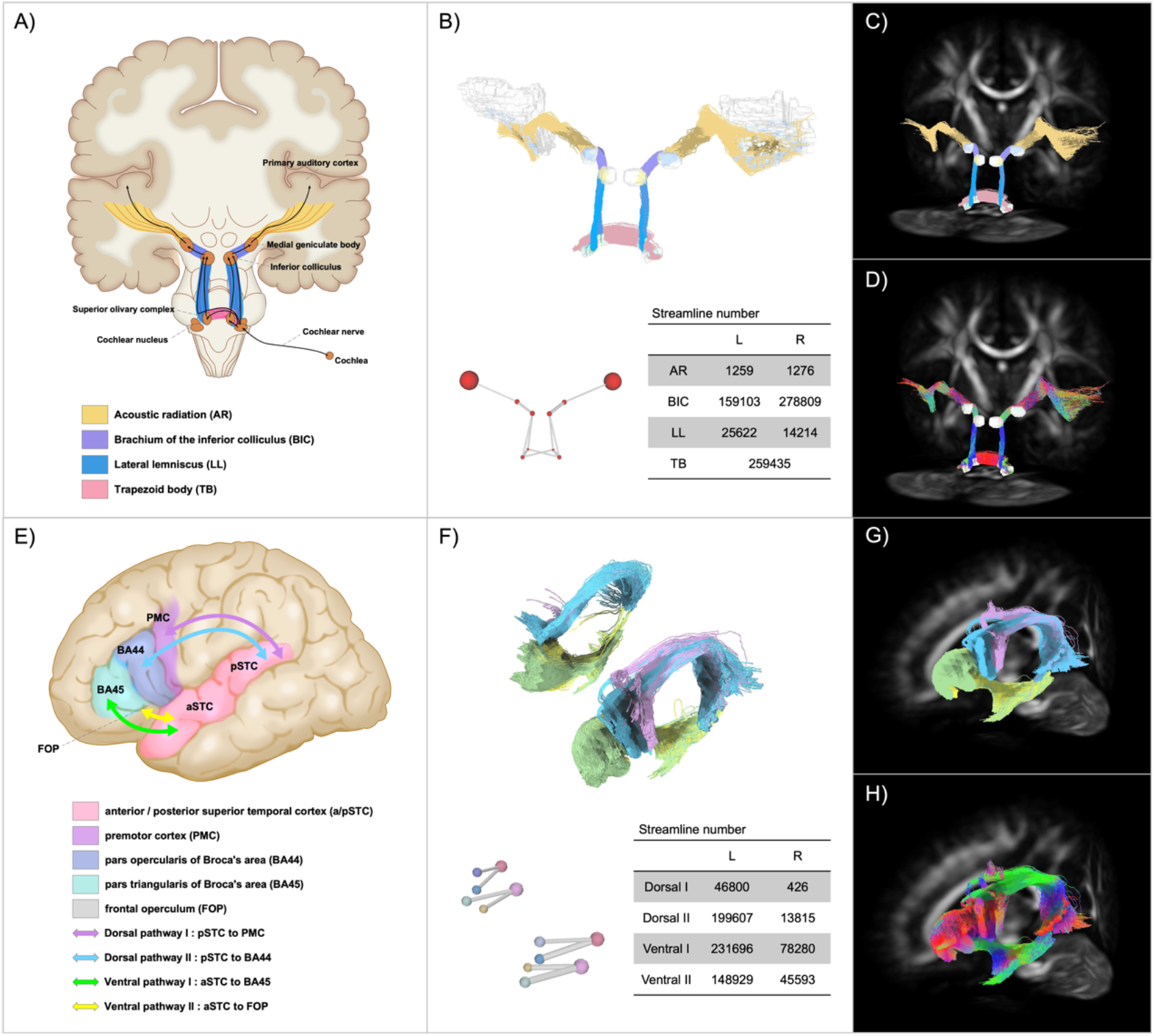
In vivo tractography of the auditory pathway and the language pathway in the study-specific template. A) and E) present schematic diagrams for the ascending auditory pathway and the language pathway, respectively. The auditory diagram was adapted from *Duvernoys Atlas of the Human Brain Stem and Cerebellum (2009)*; the language diagram was adapted from *Friederici (2017)*. B) and F) show three-dimensional reconstructions of tractography results of the central auditory pathway and the language pathway, respectively. Fibre colours refer to the subdivisions in each pathway, corresponding with the schematic colours in A) and E). The ball-and-stick diagrams represent the relative ROI size and streamline numbers in each pathway. Tractography results are also displayed in the study-specific WM FOD template in C) and G), colour coded by subdivision, and in D) and H), colour coded by direction (red: left-right; green: anterior-posterior; blue: superior-inferior). See ***Appendix -video1*** and ***Appendix -video2*** for three-dimensional animated videos of these pathways.

### Tractography of the language pathway

The language pathway was also reconstructed bilaterally, each comprising two dorsal streams and two ventral streams (see ***Figure 2***). Dorsal pathway I connects the posterior part of the superior temporal cortex (pSTC) to the premotor cortex (PMC) via the arcuate fascicle (AF) and the superior longitudinal fascicle (SLF). Dorsal pathway II connects the pSTC to the pars opercularis of Broca’s area (BA44) via the AF/SLF. Ventral pathway I connects pars triangularis of Broca’s area (BA45) and the temporal cortex via the extreme fibre capsule system (EFCS). Ventral pathway II connects the frontal operculum (FOP) and the anterior part of the STC via the uncinate fascicle (UF). The whole language pathway showed left dominance in fibre numbers.

### Children with profound SNHL exhibited fibre impairment in the central auditory pathway and the language pathway

In the central auditory pathway, FBA results demonstrated reduced FC, FD, and FDC in TB and decreased FC in LL in patients with profound SNHL (pFWE < 0.05) (see ***Figure 3A***). There was no significant difference in the “superior” part of the auditory pathway (ie. the bilateral BIC and AR). In the language pathway, only the left ventral streams showed reduced FC and only the left dorsal streams showed reduced FD, while decreased FDC was found in both the left dorsal and left ventral streams (see ***Figure 3B***). The size of impaired areas in the dorsal streams was larger than that in the ventral streams. No significant difference was found in fibre metrics of the right language pathway.

**Figure 3.**
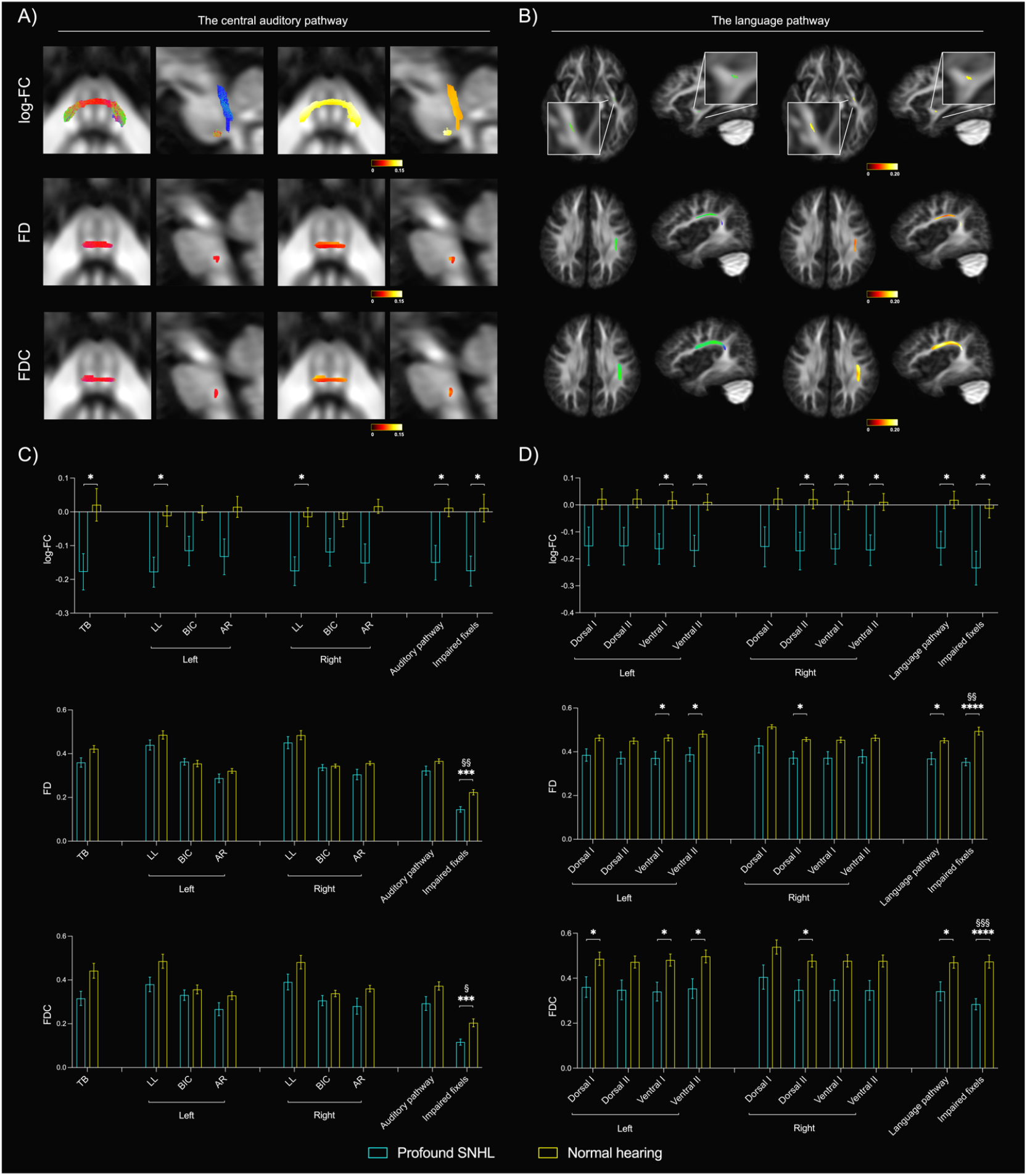
Fibre impairment of the central auditory pathway and the language pathway in children with profound SNHL. Streamlines associated with significantly reduced fibre cross-section (FC), fibre density (FD), and fibre density and cross-section (FDC) (family-wise error (FWE)-corrected p-value < 0.05) in fixel-wise comparison between patients with profound SNHL and normal hearing controls are shown for the central auditory pathway (A) and the language pathway (B). The left two columns in each panel display colour coded by direction and the right two coded by absolute values of effect size. Mean FC, FD, and FDC extracted from pathway subdivisions, entire pathways, and impaired fixels from (A) and (B) in the central auditory pathway (C) and the language pathway (D) are shown for patients with profound SNHL versus normal hearing controls. * p<0.05, ** p<0.01, *** p<0.001, **** p<0.0001; § pFWE<0.05, §§ pFWE<0.01, §§§ pFWE<0.001.

Next, we performed a tract-of-interest analysis to examine specific subdivisions in these pathways (see ***Figure 3C and 3D***). Fibre metrics of all tracts displayed a decreased trend in children with profound SNHL. In accordance with fixel-wise comparison results, only the “inferior” subdivisions of the central auditory pathway (TB and bilateral LL) had significant fibre impairments. In the language pathway, significant impairment was found bilaterally rather than only on the left side. The left ventral I, left ventral II, and right dorsal II streams exhibited all-round fibre impairment, as FC, FD, and FDC were all significantly reduced. When considering central pathways as a whole, the mean FC of the entire central auditory pathway and the mean FC, FD, and FDC of the entire language pathway were significantly reduced. After extracting the mean values of significant results from fixel-wise comparison, the mean fibre metrics of these impaired fixels demonstrated a significant, large decrease. Of all tract-of-interest comparisons, only the decrease of FD and FDC of impaired fixels in both pathways survived family-wise error (FWE) correction (pFWE<0.05).

### Peripheral auditory structure moderated the structural development of central pathways

Currently, surgical decisions are made based on visual inspection of the inner ear and cochlear nerve structure, which has limited value. In the present study, all seven children who underwent CI surgery had normal inner ear and cochlear nerve structure (see ***Table 1*** and ***Figure 4A***). All six children who underwent ABI surgery or were ABI candidates presented with inner ear malformation (IEM) and cochlear nerve deficiency (CND) (see ***Table 1*** and ***Figure 4B***). The cochlear nerve and vestibular nerve converge to form cranial nerve VIII (the vestibulocochlear nerve; cn.VIII), which travels through cerebrospinal fluid (CSF) in the cerebellopontine angle cistern and enters the brainstem. We measured the median contrast value of cranial nerve VIII (regressing out surrounding CSF median contrast values) to represent peripheral nerve tissue density (see ***Figure 4C***). One patient with absent cochlear nerve also presented with no vestibular nerve; therefore, he was absent of cn.VIII and was left out in the following statistics.

**Figure 4.**
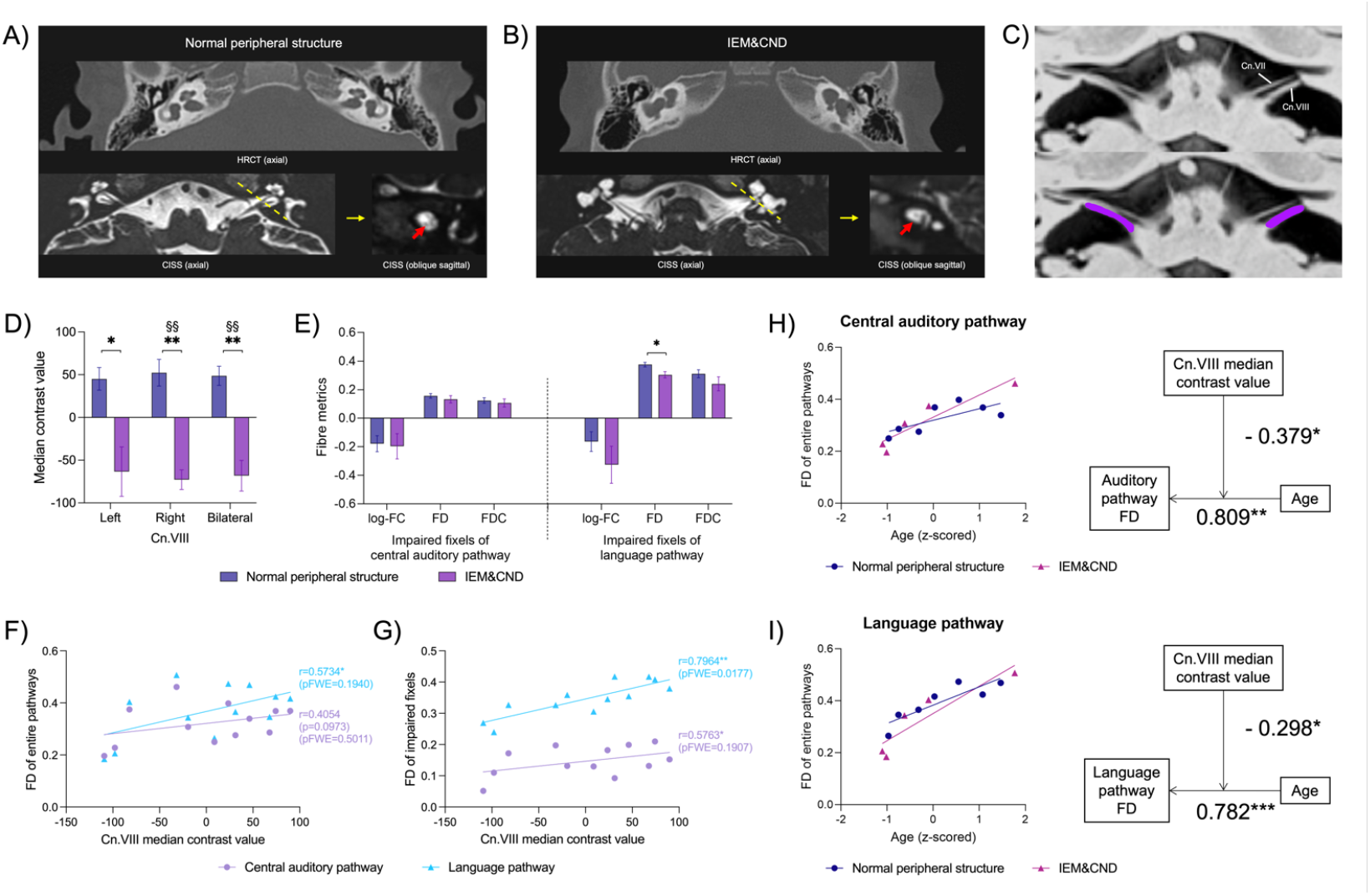
Cranial nerve VIII median contrast values and central pathway fibre metrics in profound SNHL patients with normal peripheral structure or those with IEM&CND. A) shows temporal bone HRCT and CISS sections from a patient who underwent CI surgery (male, 32 months old) and demonstrates a normal structure of the inner ear and cochlear nerve. B) presents temporal bone HRCT and CISS sections from an ABI candidate (male, 17 months old) and reveals inner ear malformation (IEM) and cochlear nerve deficiency (CND). The red arrow in (B) pointed to the missing cochlear nerve that is normally present, as shown by the red arrow in (A). C) displays inverted CISS sections (axial plane) at the pontomedullary junction from a patient with profound SNHL. Cranial nerve VIII (cn.VIII) is visualized as a hyperintense structure relative to the surrounding cerebrospinal fluid (CSF). Cn.VIII was segmented (purple) and extracted for its median contrast value, regressing out surrounding CSF median values. D) shows the cn.VIII median contrast values for patients with normal peripheral structure versus those with IEM&CND. E) presents the mean FC, FD, and FDC of deafness-associated impaired fixels (from fixel-wise comparison results between patients and controls; see ***Figure 2A, 2B***) in central pathways for patients with normal peripheral structure versus those with IEM&CND. F) displays the Pearson correlation between cn.VIII median contrast values and the mean FD of entire central pathways for patients with profound SNHL. G) illustrates the Pearson correlation between cn.VIII median contrast values and the mean FD of deafness-associated impaired fixels in central pathways for patients with profound SNHL. H) demonstrates the moderation of central auditory pathway maturation by cn.VIII median contrast values. The mean FD of the entire central auditory pathway was significantly associated with age (beta value = 0.809), and their association was negatively moderated by cn.VIII median contrast values (interaction beta value = -0.379). I) shows the moderation of language pathway maturation by cn.VIII median contrast values. The mean FD of the entire language pathway was significantly associated with age (beta value = 0.782), and their association was negatively moderated by cn.VIII median contrast values (interaction beta value = -0.298). These moderation effects are visualized as separate correlation plots of central pathway FD and age for patients with normal peripheral structure and those with IEM&CND. See ***Appendix -table 2*** and ***Appendix -table 3*** for detailed statistics of the moderation analysis. * p<0.05, ** p<0.01, *** p<0.001; § pFWE<0.05, §§ pFWE<0.01, §§§ pFWE<0.001.

**Figure 5.**
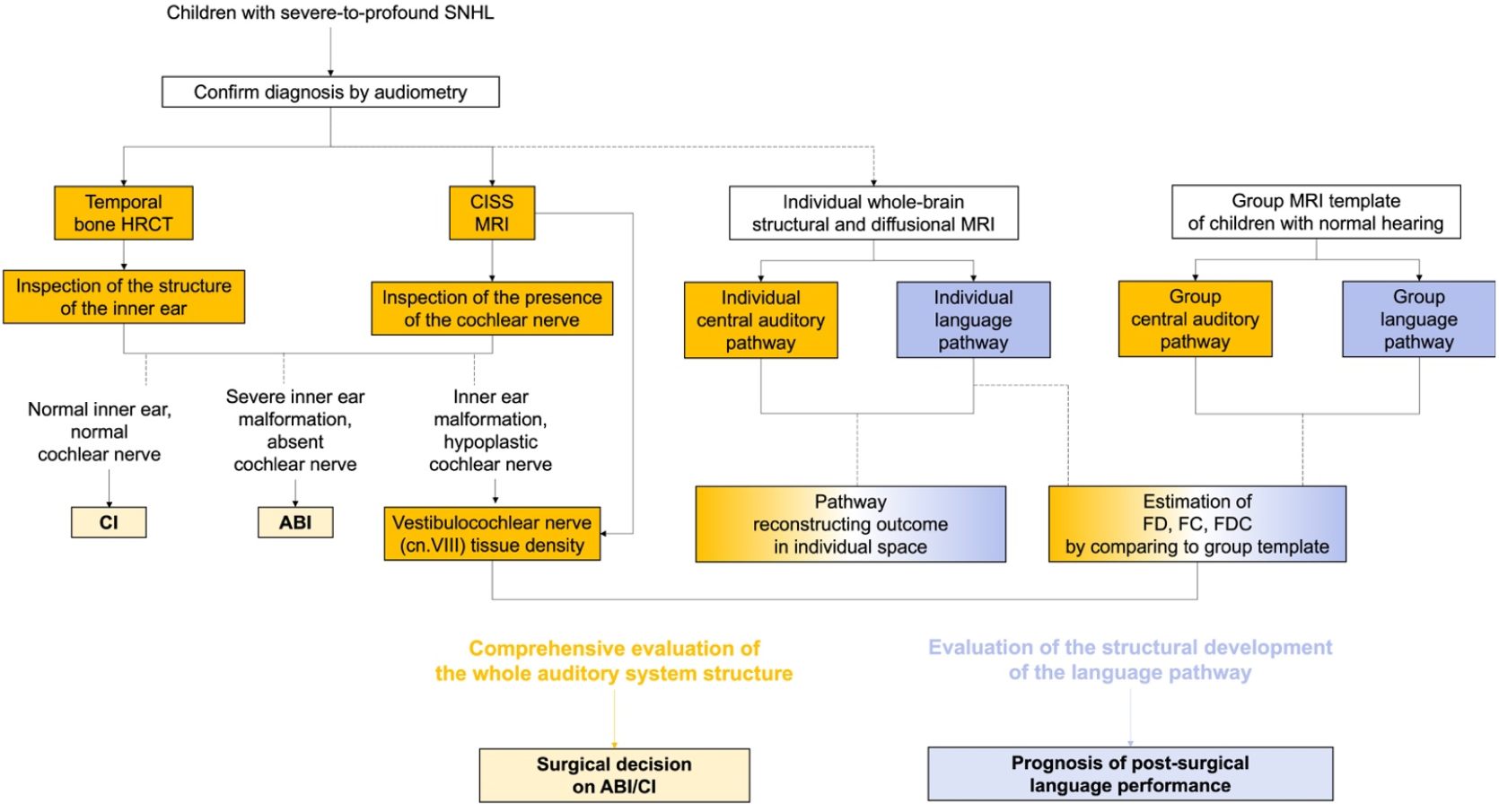
Comprehensive pre-surgical evaluation of the auditory-language network.

We divided patients with profound SNHL into a normal peripheral structure subgroup and an IEM&CND subgroup, and compared their peripheral nerve and central pathway structure. The IEM&CND subgroup had significantly lower cn.VIII median contrast values than the normal peripheral structure subgroup (pFWE<0.01; see ***Figure 4D***). Fixel-wise comparison showed no significant difference between the two subgroups in the central auditory pathway and the language pathway (pFWE>0.05). Then, we extracted mean fibre metrics of deafness-associated impaired fixels (significant regions in the fixel-wise comparison between patients with profound SNHL and normal hearing controls) and compared them between the two subgroups. All fibre metrics showed a reduced trend for the IEM&CND compared to the normal peripheral structure subgroup; only FD of deafness-associated impaired fixels in the language pathway showed a significant decrease but did not survive family-wise error (FWE) correction (see ***Figure 4E***).

To investigate the relationship between peripheral nerves and central pathways, we performed a Pearson correlation between cn.VIII median contrast values and central pathway fibre metrics. When examining mean metrics of entire pathways, FD of the language pathway, rather than FD of the central auditory pathway, was significantly correlated with cn.VIII median contrast values (r=0.5734, p<0.05, pFWE>0.05; see ***Figure 4F***). Similarly, for deafness-associated impaired fixels, the correlation between FD of the language pathway and cn.VIII median contrast values was stronger and more significant compared to the central auditory counterpart (r=0.7964 vs. r=0.5763, pFWE<0.05 vs pFWE>0.05; see ***Figure 4G***). No significant correlation was found in FC of entire pathways or deafness-associated impaired fixels. We also examined peripheral correlation with central pathway subdivisions. More correlations were found for language subdivisions than central auditory ones (see ***Appendix -table 1***).

Further, a moderation analysis was conducted to explore the specific impact of peripheral nerve structure on the maturation of central pathways over time. The temporal developmental trajectories of FD of both the entire central auditory pathway and the entire language pathway were negatively moderated by cn.VIII median contrast values (interaction beta value = -0.379 and -0.298, respectively; see ***Figure 4H and 4I, Appendix -table 2, and Appendix -table 3***). No significant moderation effect was found on the maturation of FC of central pathways by peripheral nerve structure.

## Discussion

In this study, we successfully segmented subcortical auditory regions and reconstructed the auditory and language pathways in vivo. Our findings showed decreased fibre density and fibre cross-section mainly in the inferior part of the central auditory pathway and the left language pathway. Additionally, we discovered that the correlation between language pathway fibre metrics and peripheral vestibulocochlear nerve structure is stronger and more significant than that in the central auditory pathway, and that the peripheral structure moderated the developmental trajectory of the central auditory and language pathways.

### The new pipeline for mapping the human auditory pathway with in vivo MRI

To address the issue of inability to precisely locate subcortical auditory nuclei using sound-stimulating fMRI tasks for deaf patients, we have introduced a new pipeline that only requires the acquisition and post-processing of structural and diffusional images. The cochlear nucleus appears as an angulated wedge shape along the brainstem surface when viewed from above; it is mostly located between the inferior cerebellar peduncle and cerebrospinal fluid, with a small width of up to around 2 mm (Rosahl & Rosahl, 2013). The superior olivary complex is a small cell mass that is buried deep in the brainstem, surrounded by multiple fibre bundles. These two delicate auditory nuclei are prone to partial volume effects due to insufficient resolution and a lack of contrast to differentiate them from their surroundings in in-vivo structural MR brain scans (even when using 7T MRI scanner (Sitek et al., 2019)). To address this, we have reconstructed super-resolution track density imaging (TDI) maps to provide complementary contrast for high resolution structural images, allowing the cochlear nucleus and the superior olivary complex to be delineated according to track density and track direction at an isotropic resolution of 0.5 mm. The super-resolution properties of TDI maps have been validated using in vivo and in silico data, and the anatomical contrast of TDI maps from ex vivo mouse data has been related to histology (Calamante et al., 2011; Calamante et al., 2012). TDI maps have been demonstrated to be useful in delineating substructures of the thalamus, the basal ganglia, and brainstem fibre bundle cross-sections to assist fibre tracking (Calamante et al., 2013; Kwon et al., 2021; Tang et al., 2018). Epprecht et al. found that significantly different fibre orientations can be detected in the cochlear nucleus area and in the inferior cerebellar peduncle area using diffusion tensor model (Epprecht et al., 2020), which suggests the potential role of fibre direction information in distinguishing the cochlear nucleus from adjacent structures. In the present study, the good inter-rater reliability of segmentation suggests that this method is effective for locating the subcortical auditory nuclei.

Our results showed that the best way to capture fibre features of different tracts in the auditory pathway is to track them separately and optimize tracking strategies for each part. We implemented probabilistic tractography and deliberately chose seeds and termination ROIs for each tract based on anatomical prior knowledge. MCP fibres were excluded when tracking the trapezoid body, which accounted for a large proportion of contralateral streamlines if not controlled and have yet been neglected in earlier studies. Exclusion ROIs are also essential when tracking the acoustic radiation, because the acoustic radiation crosses with several major fibre bundles that own greater FOD amplitudes in the corresponding directions. Mafei et al. found that optimal tracking parameters for the acoustic radiations were probabilistic tractography with default settings (angle threshold = 45°, step size = 1/2 * voxel size); however, the FOD amplitude threshold was not mentioned (Maffei et al., 2019). We examined a range of cutoff values and found that thresholding at 0.05 obtained optimal results for the acoustic radiations and 0.1 for subcortical auditory tracts. The tractography parameters established in the present study offer reliable recommendations for future attempts to track the auditory pathway.

### Fibre impairment pattern in the central auditory pathway of children with profound SNHL

Normally, the axonal myelination of the auditory pathway starts at the 26th foetal week, becomes definitive by the 29th week of gestation, and continuously increases in density until at least 1-year postnatal age (Moore et al., 1995). With the deprivation of hearing inputs, the central auditory pathways of congenitally deaf children displayed a brainstem-dominant fibre impairment pattern that included both microstructural impairment and macroscopic deficiency: the FD, FC, and FDC of the trapezoid body and the FC of the lateral lemnisci were significantly decreased; nevertheless, no significant difference was found in the branchium of inferior colliculus or the acoustic radiation. Current results are partly contradictory to earlier findings using the diffusion tensor model.

Huang et al. manually delineated six ROIs along the auditory pathway, including nuclei and fibre bundles, using a 14 mm^2^ square box in T2w and FA maps of children with profound hearing loss. They measured DTI-based metrics in the TB, SOC, IC, MGB, AR and white matter of Heschl’s gyrus (LL and BIC not included), and found decreased FA values in all of these six ROIs (Huang et al., 2015). Wu et al. focused on the AR and STG, demonstrating reduced FA values in both regions in children with hearing loss (Wu et al., 2016). The discrepancy between the present study and earlier findings may be caused by methodology factors. Unlike DTI metrics, FBA is capable of measuring individual fibre properties in fibre crossing areas, which is the case along almost the entire auditory pathway. Based on our new pipeline, each fibre bundle in the auditory pathway was defined by tractography separately; FBA metrics were measured in the fixel mask built on the auditory pathway so that presumable contamination from adjacent crossing fibres was excluded. Also, the Family-wise error (FWE) correction used in FBA is a more rigid multiple comparison method. As a result, we were able to make an accurate and strict examination of the fibre properties of the auditory pathway.

The trapezoid body was the most affected tract in the auditory pathway of congenital deaf children. The TB is essential to sound localization because it transfers binaural sound information to the SOC, where interaural level and time differences are compared to identify the sound source. Decreased FD and FDC in the trapezoid body indicate an immature myelination or axon loss, which may affect the efficiency of signal transmission. For patients with bilateral cochlear implants, their accuracy and sensitivity of sound localization are still worse than those of normal hearing listeners and hearing aid users (Dorman et al., 2016; Verschuur et al., 2005). One reason for this may be the absent temporal fine structure cues and limited absolute level judgements in the CI system. Another possibility is that the structural impairment of the trapezoid body plays a role in the process. If this is true, it raises the interesting question of whether auditory implants may promote structural plasticity in the trapezoid body that could enhance sound localization performance over time.

The formation of a brainstem-dominant impairment pattern warrants investigation. The absence of a significant difference in two high-level auditory pathways, the BIC and AR, may reflect that the susceptibility of fibre structures to auditory deprivation decreases in a bottom-up fashion along the pathway, or that upper pathways in the thalamus and cerebrum may have already undergone cross-modal plasticity so that fibres are still structurally intact but subserve other functions. It has been proposed that the auditory cortex might reorganize to mediate other functions such as vision. Auditory areas in the superior temporal sulcus show greater recruitment in deaf individuals than in hearing individuals when processing visual, tactile or signed stimuli (Bavelier et al., 2006). However, it is unclear that to what extent the primary auditory cortex may be affected by cross-modal plasticity and whether such cortical plasticity would affect downstream auditory nuclei and in-between fibre bundles. Further studies incorporating structural and functional features of the auditory pathway in larger samples are required to shed light on these questions.

### The structural development of the language pathway in children with profound SNHL

After CI and/or ABI implantation, children were able to achieve good hearing sensation, but their performance in speech recognition and production was poorer and varied (Sennaroğlu et al., 2016). One of the most important tasks in this field is identifying the factors that contribute to language outcome after implantation. In this study, we focused on the structural properties of the language pathway and found that patients with profound SNHL had an all-streams-affected fibre impairment that was left-dominant.

The sub-structures of the language pathway are closely connected to language functions. The coordination between BA44 and pSTC subserves syntactic computation, while the ventral streams that connect BA45 and FOP with the temporal cortex support lexical-semantic comprehension. The pSTC integrates syntactic and semantic information for comprehension at the sentence level and is also connected to the premotor cortex as a peripheral sensorimotor interface system (Friederici et al., 2017). The maturational status of these tracts has been found to be associated with behavioural performance. For example, the tract targeting BA44 is highly predictive of behavioural performance on processing hierarchically complex sentences (Skeide et al., 2016). In this study, these four streams were separately reconstructed using probabilistic tractography. FBA results showed that the dorsal streams mainly suffered from reduced fibre density, while the ventral streams mainly exhibited decreased fibre cross-section in children with profound SNHL. The combined metric FDC was more sensitive in detecting fibre impairment in both dorsal and ventral streams. These results demonstrate that deprived auditory inputs have a damaging effect on the structural development of all language streams that serve different functions, which may indicate that congenitally deaf children suffer from an overall underdeveloped language capacity in semantics, syntax, and sensorimotor integration, rather than just struggling with oral communication.

### Moderation of central pathway maturation by peripheral auditory structure

Patients with normal peripheral auditory structure and those with IEM&CND had significantly different cn.VIII median contrast values (which has been shown to represent nerve tissue density (Harris et al., 2021)), but few differences in fibre density and fibre cross-section of the central auditory pathway and language pathway. However, several central pathways were positively correlated with cn.VIII median contrast values, including FC of bilateral BIC and LL, FC of two left ventral streams, and FD of bilateral dorsal streams and left ventral streams. Although all patients included in the study were diagnosed with bilaterally profound SNHL, several patients with normal peripheral structure had late-onset or progressive hearing loss that may have allowed a small amount of peripheral auditory inputs to stimulate central development before complete deafness. In contrast, all patients with cochlear nerve deficiency failed newborn hearing screening and remained deaf since. This phenomenon may partly explain the correlation between peripheral nerve tissue density and central fibre metrics.

Language pathway fibre metrics showed stronger and more significant correlations with peripheral nerve tissue density than central auditory ones in the case of entire pathways, pathway subdivisions, and deafness-associated impaired fixels. These findings may deviate from expectations since the central auditory pathway is directly connected to the peripheral cochlear nerve both structurally and functionally. For profound SNHL patients with normal-appearing cochlear nerves and inner ear structure, their primary pathology is usually confined to the cochlea and occurs at the cellular level. In contrast, non-syndromic profound SNHL patients with cochlear nerve deficiency, who have lower cn.VIII tissue density and accompanying bony inner ear malformations, are considered to have more severe genetic abnormalities. It is still unclear whether these patients have anomalies in the central auditory pathway caused by the same genetic factors that lead to peripheral malformation or have underdeveloped central auditory structures due to auditory deprivation after peripheral dysfunction. The weak relationship between deafness-associated impaired fixels in the central auditory pathway and cn.VIII tissue density makes the former hypothesis less likely.

On the other hand, the significant association between FD/FC of deafness-associated impaired language fixels and cn.VIII tissue density suggests that the language pathway is more sensitive to peripheral auditory condition compared to the central auditory pathway. The language pathway, as the substrate for higher cognitive functions, develops in a way that is highly dependent on external inputs and interaction with the environment (Kuhl & Rivera-Gaxiola, 2008). More specifically, acquiring proper inputs during a sensitive period is essential for the maturation of the language pathway. This structural developmental feature corresponds with behavioural results from follow-up studies on CI recipients of different age at implantation. Chen et al. compared performances of children implanted before and after two years old and found that the younger the age of bilateral cochlear implant surgery, the higher developmental quotient of language, social skills, and adaptability the child could achieve after two years (Chen et al., 2022). Houston et al. studied an earlier demarcation timepoint at one year old and found that children implanted during the first year of life had better vocabulary outcomes than children implanted during the second year of life; however, earlier implanted children did not show better speech perception outcomes than later implanted children. They suggested that there is an earlier sensitive period for developing the ability to learn words than for central auditory development (Houston & Miyamoto, 2010).

To understand the specific impact of peripheral structure on the developmental trajectory of central pathways, we conducted a moderation analysis. Cn.VIII tissue density had a significant negative moderation effect on the relationship between auditory pathway FD and age, and that between language pathway FD and age. In other words, the lower the cn.VIII tissue density, the closer the relationship between fibre density of central pathways and age. It is worth noting that cn.VIII tissue density only moderated FD maturation; there was no significant moderation effect on FC maturation. These results may imply that the macroscopic developmental trajectory of central pathways is similar among patients with various peripheral conditions, while at the microscopic level, peripheral nerve deficiency may lead to a delayed and slowed axon generation or myelination in central pathways, which may reflect as an increased association of central fibre density with age in the participants ranging from six months to six years old. Taken together, to prevent aberrant structural development in the auditory-language network in the absence of hearing, auditory interventions should be implemented as early as possible. Special attention should be paid to patients with cochlear nerve deficiency. Early interventions would provide children with timely auditory inputs during the critical period of language development and lead to better long-term outcomes.

### Comprehensive pre-surgical evaluation of the auditory-language network

Our work fills in the gap of quantitative examination of the central auditory structural development and provides an individualized, comprehensive pre-surgical evaluation of the auditory-language network to assist surgical decisions and estimate prognosis. By optimizing the CISS-based estimation of cn.VIII tissue density, we allow for a more accurate quantitative measurement of peripheral nerve structure, which is essential for characterizing the extent of cochlear nerve deficiency. By reconstructing the entire central auditory pathway at the individual level, we can measure fibre density and fibre cross-sections to estimate the structural maturation and integrity of the central auditory pathway, which may imply the efficiency of auditory information transmission after implantation. Combining the findings from HRCT, CISS MRI, and reconstruction of the central auditory pathway enables a comprehensive view from the cochlea to the auditory cortex to help make surgical decisions, such as choices between CI and ABI and the side of implantation. Additionally, by measuring the fibre properties of the language pathway, we were able to evaluate the structural development of the language network, which shows great potential for estimating post-implant language performance. Investigating the maturation of specific streams in the language pathway also offers insights into the corresponding language subfunctions (syntax, semantics, or sensorimotor integration) that should be emphasized during post-implant language training.

### Strengths and limitations

The strengths of the present study include: (1) the first comprehensive in vivo reconstruction of the central auditory pathway independent of using auditory stimuli; (2) the inclusion of children with inner ear malformations and cochlear nerve deficiency in addition to those with normal inner ear structures to provide a complete picture of congenitally deaf children; (3) a precise inspection of fibre structures of the auditory pathway, the language pathway, and their subdivisions using a fixel-based approach that address crossing fibre issues. We also acknowledge some limitations to this study. The sample size is relatively small, and follow-up behavioural evaluations of hearing and speech capacity after CI and ABI surgeries would provide more information on the prognostic value of neuroimaging findings.

### Conclusions

In conclusion, this study introduced a new pipeline for in vivo reconstruction of the central auditory pathway, found both microscopic and macroscopic fibre impairment in specific auditory and language tracts, and discovered a negative moderation effect of peripheral auditory structure on central pathway maturation. We provided structural evidence supporting the necessity of early auditory intervention and established a promising comprehensive pre-surgical evaluation of the auditory-language network for children with severe-to-profound SNHL to assist with surgical planning and prognosis.

## Methods

### Study sample

Twenty-three children aged under six years old, including thirteen patients with bilateral profound congenital sensorineural hearing loss and ten controls matched on age and sex, were included.

The patients met the following criteria: (1) diagnosis of bilateral profound sensorineural hearing loss: click auditory brainstem response (ABR) threshold > 95 dB nHL, or pure-tone average (PTA, 0.5-4k Hz) threshold > 95 dB HL (Lin et al., 2011); and (2) non-syndromic hearing loss (no association with any other systemic manifestations). The controls had normal hearing in both ears. Exclusion criteria for all volunteers were as follows: (1) neurological disease (epilepsy, brain tumour, etc.) or history of head trauma; (2) psychiatric disorders (autism spectrum disorder, etc.); (3) history of ototoxic drug use; and (4) metal implants and other contraindications to MRI safety.

The study protocol was approved (SH9H-2021-T449-1) by the Ethics Committee of Shanghai Jiao Tong University School of Medicine Affiliated Ninth People’s Hospital (Shanghai, China), and all enrolled subjects had informed consent provided by parent/guardian.

### Clinical data

Age, sex, gestational weeks, birth weight, and a thorough medical history of pregnancy and hearing condition were recorded.

Pure tone audiometry at 0.5k, 1k, 2k, 4k, and 8k Hz (for children who were able to cooperate) and click ABR were examined to evaluate hearing levels (AC40 and Eclipse, Intercoustics, Middelfart, Denmark).

Nine out of thirteen patients underwent auditory implantation (two ABI recipients and seven CI recipients). Post-implant outcomes of hearing and language development were assessed, during and 1, 3, 6, 12, 18, and 24 months after device activation, using pure tone audiometry and the following scales: (1) Categories of Auditory Performance (CAP) (Archbold et al., 1995); (2) Speech Intelligibility of Rating (SIR) (Allen et al., 1998); (3) Infant-toddler Meaningful Auditory Integration Scale (IT-MAIS); and (4) Meaningful Use of Speech Scale (MUSS) (Zhong et al., 2017). The follow-up evaluation is in progress and the results will be reported in our subsequent studies.

### CT, MRI acquisition and quality assessment

A temporal bone high-resolution CT (sections of 0.5 mm in thickness) and a three-dimensional (3D) constructive interference in steady state (CISS) MRI scan (3-Tesla MAGNETOM Vida, Siemens Healthcare, Erlangen, Germany) were acquired to inspect the structure of the inner ear and the cochlear nerve of patients. Two experienced neuroradiologists assessed the presence and type of inner ear malformation and cochlear nerve deficiency based on Sennaroğlu classification criteria (Sennaroğlu & Bajin, 2017).

Whole brain MRI was carried out on a 3-Tesla Siemens MAGNETOM Vida scanner (Siemens Healthcare, Erlangen, Germany) using a 64-channel head coil. To reduce motion artifacts, patients received sedation by oral intake of chloral hydrate for the MRI scan under supervision. Earplugs and sound-attenuating foam were used to decrease the noise of the scanner. Sandbags were placed on the scanner table to help attenuate vibrations during scanning. The lights in the scanner room were turned off, and blankets were wrapped around the children to create a suitable sleeping environment. Monitoring throughout the scanning session included pulse oximetry, respiration, and close observation by medical staff.

The acquisition protocol was adapted from the Developing Human Connectome Project (DHCP) that was optimized for the properties of the developing brain (Edwards et al., 2022). All children underwent a T1-weighted anatomical brain scan (3D MPRAGE sequence with a spatial resolution of 0.8 mm isotropic, matrix = 320 × 320, field of view (FOV) = 256 × 256 mm^2^, and TR/TE = 2400/2.38 ms) and whole brain diffusion imaging (dMRI). The scanning protocol consisted of three diffusion shells (b-values of 400, 1000, and 2600 s/mm^2^) with 32, 44, and 64 diffusion-weighted directions each and ten b0 volumes using PA phase encoding. Additionally, two b0 images in AP phase encoding were scanned for TOPUP distortion correction. The EPI readout used SMS factor of 4, Grappa acceleration factor of 2, partial Fourier factor of 6/8, isotropic resolution of 2.0 mm, axial slices of 72, matrix of 105 × 105, FOV of 210 × 210 mm^2^, and TR/TE of 2900/95 ms.

Each patient’s MRI was transferred to a DICOM workstation during acquisition to review any clinical or research-relevant incidental findings. Then, insufficient coverage, excessive motion, and/or ghosting were visually assessed. If any image failed the visual quality assessment, the MR technicians would decide whether to re-scan the sequence according to the child’s condition.

### Cranial nerve VIII measurement

The measurement of cranial nerve VIII tissue density was adapted from Harris et al.’s protocol (Harris et al., 2021). Specifically, the contrast of CISS images was inverted to visualize cranial nerve VIII as a hyperintense structure relative to the surrounding CSF. Cn.VIII was segmented on each axial section using ITK-SNAP (Yushkevich et al., 2006) by two independent raters (inter-rater DICE coefficient = 0.977±0.010). The median contrast value of cn.VIII was calculated across sections. The median contrast values of adjacent CSF were also collected and regressed out to control contrast differences across individuals due to scanner heating and motion artifact.

### Diffusional MRI processing

The state-of-the-art fixel-based analysis (FBA) pipeline (Dhollander et al., 2021) was implemented to process diffusional data using MRtrix3 (Tournier et al., 2019). Specifically, after standard preprocessing steps (including denoising, Gibbs ringing correction, eddy-current and motion correction, bias field correction, and intensity normalisation), response functions for WM, GM and CSF were estimated from the data themselves. The diffusional images were then upsampled to 1.25 mm isotropic voxels for subsequent better estimation of fibre orientation distribution (FOD). We used multi-shell multi-tissue constrained spherical deconvolution (msmt-CSD) to obtain WM-like FOD as well as GM-like and CSF-like counterparts in all voxels (Dhollander & Connelly, 2016). A study-specific WM FOD template was created using the WM FOD images from all ten controls. Finally, study-specific auditory and language pathways were generated from this template and filtered to reduce reconstruction bias (Smith et al., 2015). These generated tractograms were then converted to fixel masks, allowing for fixel-based analysis of specific tracts.

### Reconstruction of the human auditory pathway

#### Overview

We generated directionally encoded colour track density imaging (DEC-TDI) maps from whole-brain tractography to obtain high spatial resolution images of the white matter. These DEC-TDI maps and T1-weighted images provided complementary information and enhanced anatomical contrast for subsequent manual segmentation of subcortical auditory nuclei. The primary auditory cortex was extracted from the Human Brainnetome Atlas (Fan et al., 2016) and co-registered to diffusional space. Finally, the auditory pathway was tracked based on anatomical prior knowledge and visualised using 3D volume rendering. This process was performed at both the group-average and individual level.

#### Track density imaging

The DEC short-tracks TDI (stTDI) map method (Calamante et al., 2012) was used to obtain better directional information compared to the standard DEC-TDI pipeline (Calamante et al., 2010), particularly in low-intensity structures such as brainstem nuclei that we were interested in. Whole brain probabilistic tractography was constrained to short tracks by setting the maximum length of each track to 20 mm (corresponding to 10 acquired voxels). We generated 40 million short tracks for each dataset using the iFOD2 algorithm (Tournier et al., 2010) by randomly seeding throughout the brain with the following parameters: angle threshold = 45°, minimum length = 4 mm, maximum length = 20 mm, cutoff value = 0.1. We then constructed the super-resolution TDI maps with a 0.5-mm isotropic grid size by calculating the number of tracks in each element of the grid. The colour-coding values were obtained by averaging the colours of all the streamline segments contained within each grid element, thereby indicating the local fibre orientation. (Green represents anterior-to-posterior, blue represents superior-to-inferior, and red represents left-to-right.)

#### Image registration

At the group level, a study-specific T1-weighted brain template was created using the T1-weighted images from all ten controls using antsMultivariateTemplateConstruction2.sh in ANTs (Avants et al., 2011) and transformed to the study-specific DEC-TDI space. At the individual level, T1-weighted images were transformed to each individual’s DEC-TDI space. Therefore, manual segmentation can be carried out in T1-weighted and DEC-TDI with a 0.5mm isotropic resolution.

#### Manual segmentation of subcortical auditory regions

Subcortical auditory regions were segmented based on anatomical observations in histology studies (Moore, 1987; Rosahl & Rosahl, 2013; Winer, 1984) as well as earlier attempts to delineate some of these structures via MRI in vivo (García-Gomar et al., 2019; Sitek et al., 2019). Two raters independently segmented the auditory nuclei using the mrview toolbox in MRtrix3 (Tournier et al., 2019). Only the overlap areas between the two raters’ segmentations were retained in the following analysis (inter-rater DICE coefficient: CN, 0.798; SOC, 0.690; IC, 0.829; MGB, 0.795).

##### Cochlea nucleus (CN)

The cochlea nucleus is located on the brainstem surface at the pontomedullary junction, where auditory nerve axons enter and terminate. The cochlear nucleus is elongated and curved from ventrolateral to dorsomedial. The ventral and dorsal portions of the cochlear nucleus can be distinguished by histological cytoarchitectonic properties, although approximately 10% of their shared borders remain a grey zone (Rosahl & Rosahl, 2013). In a horizontal view, the cochlear nucleus borders the inferior cerebellar peduncle (ICP) medially; its anterior half extends laterally along the posterior edge of the middle cerebellar peduncle (MCP) (Moore & Osen, 1979; Terr & Edgerton, 1985).

On T1-weighted images, the cochlear nucleus can be located at the pontomedullary junction where cranial nerve VIII enters the brainstem and is roughly delineated along the brainstem surface from ventrolateral to dorsomedial. In DEC-TDI maps, the cochlear nucleus is distinguished from its medial neighbour, the ICP, by its clear blue border, as the ICP travels mainly in the rostrocaudal direction (Epprecht et al., 2020). The cochlear nucleus, on the other hand, is a mixture of cell bodies and axons that travel from ventrolateral to dorsomedial, resulting in either low intensity areas (where cells dominate) or green colour areas (where axons dominate).

##### Superior olivary complex (SOC)

The superior olivary complex is a group of cells located in the pons, a short distance medial and rostral to the cochlear nucleus. The SOC is composed of a laminar medial nucleus (which extends about 4 mm rostrocaudally) and a small lateral nucleus; the entire complex is enclosed by a capsule of rostrally-directed axons of the ascending auditory pathway (Moore, 1987; Strominger & Hurwitz, 1976).

The SOC is not distinguishable on T1-weighted images. In DEC-TDI maps, the SOC appears as a hypointense area surrounded by hyperintense fibres in the horizontal view: medially, the medial lemnisci; ventrally, the trapezoid body; and laterally, the MCP. The SOC was delineated from the same axial plane as the rostral-most extent of the ventral cochlear nucleus, extending about 4 mm rostrally (García-Gomar et al., 2019).

##### Inferior colliculus (IC)

The inferior colliculus is easy to locate as the two inferior spherical structures of the corpora quadrigemina in the dorsal midbrain (Mansour et al., 2019). On T1-weighted images, the IC is distinguished from its medially adjacent structure, the periaqueductal grey matter, by demonstrating more intense T1 signals in the horizontal view. However, in DEC-TDI maps, most of the signal of the IC is lost, possibly due to distortion from tissue/air interface.

##### Medial geniculate body (MGB)

The medial geniculate body is located in the ventromedial thalamus. On T1-weighted images, the MGB is identified as an oval-shaped hypointense eminence that is medial to the lateral geniculate body and lateral to the superior colliculus (Winer, 1984). In the horizontal view of DEC-TDI maps, the MGB is restricted ventrolaterally by the blue areas of the corticospinal tract.

#### Tractography

Subcortical auditory regions were segmented manually as described above. The primary auditory cortex (A41/42, TE1.0 and TE1.2) was extracted from the Human Brainnetome Atlas (Fan et al., 2016) in the MNI space and co-registered to the WM FOD space by a rigid, affine, and nonlinear transformation using ANTs (Avants et al., 2011). Probabilistic tractography of each major tract in the auditory pathway was performed in the WM FOD space using iFOD2 algorithm with optimized parameters. For cortical tracts (acoustic radiations), we set a 0.05 cutoff value and 80 mm maximum length; for subcortical tracts, the cutoff value was 0.1 and the maximum length was set to 200 mm. Other parameters were kept the same across all tracts: 10,000 seeds per voxel, angle threshold = 45°, and minimum length = 4 mm. A mask of brainstem and thalamus was created semi-automatically using ITK-SNAP (Yushkevich et al., 2006) to constrain tracking of the subcortical auditory pathway; a centre ROI in the sagittal plane was used as exclusion for ipsilateral tracking.

##### Trapezoid body (TB) and lateral lemnisci (LL)

The neurons in the cochlear nucleus receive nerve innervation from the cochlea and project to the inferior colliculus both directly and indirectly. The superior olivary complex is the major relay station and receives axons from mostly the contralateral, partly the ipsilateral cochlear nucleus; the contralateral dominance remains in the following ascending pathway (Moore, 1987). The ascending axons below the level of the SOC form the trapezoid body; axons above the level of the SOC form the lateral lemniscus (Moore et al., 1995). The TB travels horizontally in the inferior pons and comprises many crossing fibres. The lateral lemniscus runs rostrally and dorsally to the inferior colliculus and is located on the lateral side of the brainstem superficially.

Tracking of the TB and LL was carried out by seeding from each cochlear nucleus to the SOC and the IC in both sides, using middle cerebellar peduncle (MCP) as an exclusion ROI for anatomical constraints, and seeding from each SOC to the IC in both sides.

##### Brachium of inferior colliculus (BIC)

All ascending projections from the auditory brainstem to the thalamus are carried in the brachium of the inferior colliculus (Moore, 1987). Tracking of the BIC was performed by seeding from the IC to the ipsilateral MGB. There are also commissural pathways between the bilateral IC; however, the commissure of IC is difficult to trace via tractography due to signal loss near the tissue/air interface.

##### Acoustic radiation (AR)

The acoustic radiation is the final stream that links the subcortical auditory pathway to the auditory cortex (Rademacher et al., 2002). The AR crosses with or travels near many major fibre bundles on its way to the auditory cortex, including the corticospinal tract (CST), the arcuate fasciculus (AF), the inferior fronto-occipital fasciculus (IFOF), the middle longitudinal fasciculus (MLF), the inferior fasciculus (ILF), and the optic radiation (OR). We manually delineated major cross-sections of the tracts mentioned above as exclusion ROIs. Then, we tracked the AR by seeding from the MGB, terminating in the ipsilateral primary auditory cortex, and excluding the adjacent tracts for anatomical constraint.

#### Visualization

Streamlines were transformed into the trk format in Python via the Nibabel package (https://nipy.org/nibabel/) and visualized in DSI Studio (https://dsi-studio.labsolver.org/).

### Reconstruction of the language pathway

Language ROIs were also extracted from the Human Brainnetome Atlas (Fan et al., 2016) for its finer subdivision in the temporal and frontal cortex, and co-registered from the MNI152 T1 space (Fonov et al., 2009) to the WM FOD space using rigid, affine, and non-linear transformation with ANTs (Avants et al., 2011). The anterior superior temporal cortex (aSTC) was defined as the combination of A22r, A38l, and aSTS; the posterior superior temporal cortex (pSTC) was segmented by combining A22c, rpSTS, and cpSTS. The frontal areas were also extracted: the pars opercularis of Broca’s area (BA44), pars triangularis of Broca’s area (BA45), the frontal operculum (FOP), and the premotor cortex (PMC).

Probabilistic tractography was performed in the WM FOD space. Two dorsal streams of the language pathway were seeded from BA44 or PMC and terminated in pSTC; two ventral streams were seeded from BA45 or FOP and terminated in aSTC. Parameters included: iFOD2 algorithm, 10,000 seeds per voxel, angle threshold = 45°, cutoff value = 0.1, minimum length = 4 mm, maximum length = 200 mm. Tracts were visualized in the same way as the auditory pathway.

### Fixel-based metrics

In the FBA framework, a ‘fixel’ refers to a ‘fibre population within a voxel’, allowing for the measurement of WM metrics for individual fibres crossing in the same voxel. Fibre density (FD), fibre cross-section (FC), and fibre density and cross-section (FDC) were calculated for the study-specific auditory pathway and the study-specific language pathway. FD values are approximately proportional to total intra-axonal volume and measure WM microstructure, while FC estimates macroscopic differences by using information from individual subject warps to the study-specific template. The combined FDC measure enables a more sensitive assessment of fixel-wise effects (Dhollander et al., 2021).

### Statistical analysis

Fixel-wise comparison of FC, FD, and FDC between groups was conducted using the connectivity-based fixel enhancement method (Raffelt et al., 2015). Tract-wise analysis was performed by extracting the mean fibre metrics of tracts-of-interest and comparing them between groups using PALM (Winkler et al., 2014) in MATLAB. For both comparisons, age and sex were controlled, and family-wise-corrected (FWE) p-values were obtained via permutation testing. Pearson correlation between cn.VIII median contrast values and central pathway fibre metrics were conducted using PALM in MATLAB. Moderation analysis was performed using SPSS.

## Appendix

**Appendix —table 1.**
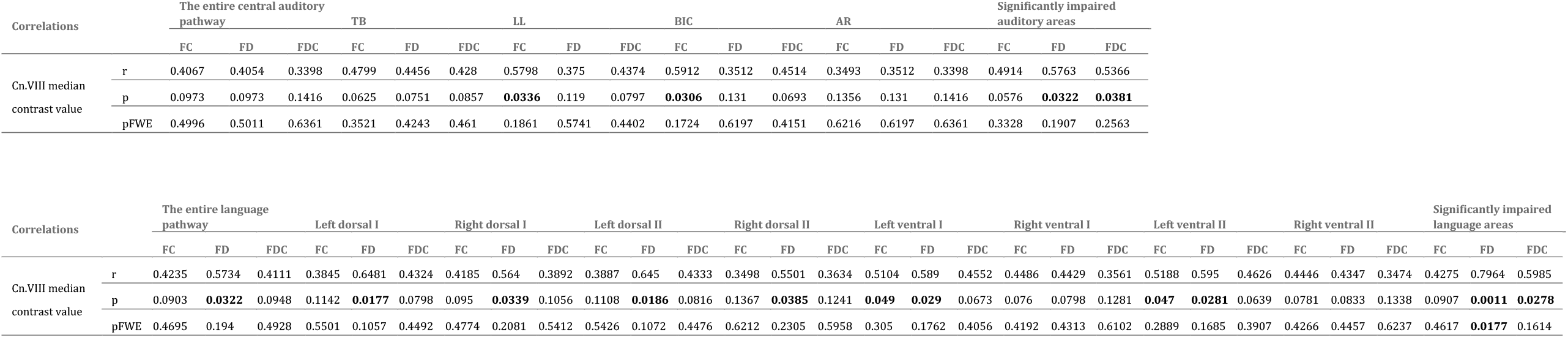
Pearson Correlation between cn.VIII median contrast values and fibre metrics of central pathways.

| Correlations |  | The entire central auditory pathway |  |  | TB |  |  | LL |  |  | BIC |  |  | AR |  |  | Significantly impaired auditory areas |  |  |
| --- | --- | --- | --- | --- | --- | --- | --- | --- | --- | --- | --- | --- | --- | --- | --- | --- | --- | --- | --- |
|  |  | FC | FD | FDC | FC | FD | FDC | FC | FD | FDC | FC | FD | FDC | FC | FD | FDC | FC | FD | FDC |
| Cn.VIII median contrast value | r | 0.4067 | 0.4054 | 0.3398 | 0.4799 | 0.4456 | 0.428 | 0.5798 | 0.375 | 0.4374 | 0.5912 | 0.3512 | 0.4514 | 0.3493 | 0.3512 | 0.3398 | 0.4914 | 0.5763 | 0.5366 |
|  | p | 0.0973 | 0.0973 | 0.1416 | 0.0625 | 0.0751 | 0.0857 | **0.0336** | 0.119 | 0.0797 | **0.0306** | 0.131 | 0.0693 | 0.1356 | 0.131 | 0.1416 | 0.0576 | **0.0322** | **0.0381** |
|  | pFWE | 0.4996 | 0.5011 | 0.6361 | 0.3521 | 0.4243 | 0.461 | 0.1861 | 0.5741 | 0.4402 | 0.1724 | 0.6197 | 0.4151 | 0.6216 | 0.6197 | 0.6361 | 0.3328 | 0.1907 | 0.2563 |

| Correlations |  | The entire language pathway |  |  | Left dorsal I |  |  | Right dorsal I |  |  | Left dorsal II |  |  | Right dorsal II |  |  | Left ventral I |  |  | Right ventral I |  |  | Left ventral II |  |  | Right ventral II |  |  | Significantly impaired language areas |  |  |
| --- | --- | --- | --- | --- | --- | --- | --- | --- | --- | --- | --- | --- | --- | --- | --- | --- | --- | --- | --- | --- | --- | --- | --- | --- | --- | --- | --- | --- | --- | --- | --- |
|  |  | FC | FD | FDC | FC | FD | FDC | FC | FD | FDC | FC | FD | FDC | FC | FD | FDC | FC | FD | FDC | FC | FD | FDC | FC | FD | FDC | FC | FD | FDC | FC | FD | FDC |
| Cn.VIII median contrast value | r | 0.4235 | 0.5734 | 0.4111 | 0.3845 | 0.6481 | 0.4324 | 0.4185 | 0.564 | 0.3892 | 0.3887 | 0.645 | 0.4333 | 0.3498 | 0.5501 | 0.3634 | 0.5104 | 0.589 | 0.4552 | 0.4486 | 0.4429 | 0.3561 | 0.5188 | 0.595 | 0.4626 | 0.4446 | 0.4347 | 0.3474 | 0.4275 | 0.7964 | 0.5985 |
|  | p | 0.0903 | **0.0322** | 0.0948 | 0.1142 | **0.0177** | 0.0798 | 0.095 | **0.0339** | 0.1056 | 0.1108 | **0.0186** | 0.0816 | 0.1367 | **0.0385** | 0.1241 | **0.049** | **0.029** | 0.0673 | 0.076 | 0.0798 | 0.1281 | **0.047** | **0.0281** | 0.0639 | 0.0781 | 0.0833 | 0.1338 | 0.0907 | **0.0011** | **0.0278** |
|  | pFWE | 0.4695 | 0.194 | 0.4928 | 0.5501 | 0.1057 | 0.4492 | 0.4774 | 0.2081 | 0.5412 | 0.5426 | 0.1072 | 0.4476 | 0.6212 | 0.2305 | 0.5958 | 0.305 | 0.1762 | 0.4056 | 0.4192 | 0.4313 | 0.6102 | 0.2889 | 0.1685 | 0.3907 | 0.4266 | 0.4457 | 0.6237 | 0.4617 | **0.0177** | 0.1614 |

**Appendix —table 2.**
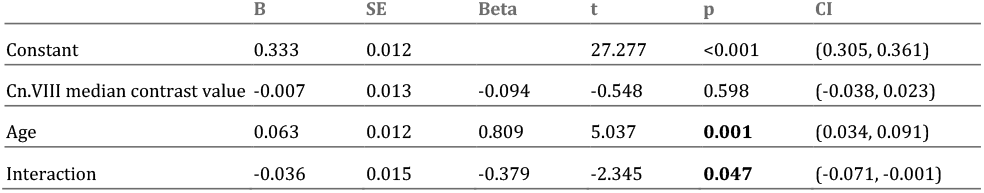
Moderation analysis of the effect of cn.VIII median contrast values on the relationship between auditory pathway FD and age.

|  | B | SE | Beta | t | p | CI |
| --- | --- | --- | --- | --- | --- | --- |
| Constant | 0.333 | 0.012 |  | 27.277 | <0.001 | (0.305, 0.361) |
| Cn.VIII median contrast value | -0.007 | 0.013 | -0.094 | -0.548 | 0.598 | (-0.038, 0.023) |
| Age | 0.063 | 0.012 | 0.809 | 5.037 | **0.001** | (0.034, 0.091) |
| Interaction | -0.036 | 0.015 | -0.379 | -2.345 | **0.047** | (-0.071, -0.001) |

**Appendix —table 3.** Moderation analysis of the effect of cn.VIII median contrast values on the relationship between language pathway FD and age.

|  | B | SE | Beta | t | p | CI |
| --- | --- | --- | --- | --- | --- | --- |
| Constant | 0.38 | 0.012 |  | 31.914 | <0.001 | (0.352, 0.407) |
| Cn.VIII median contrast value | 0.015 | 0.013 | 0.14 | 1.123 | 0.294 | (-0.015, 0.044) |
| Age | 0.081 | 0.012 | 0.782 | 6.709 | **<0.001** | (0.053, 0.109) |
| Interaction | -0.038 | 0.015 | -0.298 | -2.546 | **0.034** | (-0.072, -0.004) |

## Acknowledgements

This research was funded by Shanghai Municipal Science and Technology Commission Major Basic Research (2018SHZDZX05), Shanghai Clinical Key Specialty Construction (shslczdzk00802), Shanghai Jiao Tong University School of Medicine - High-level Local University Construction Project, Shanghai Jiao Tong University School of Medicine Collaborative Innovation Project for Translational Medicine (TM202011), Shanghai Key Laboratory for Otolaryngological Diseases Translational Medicine (14DZ2260300), and Shanghai Shen-kang Hospital Development Centre Emerging Frontier Project (SHDC12020105). The funding sources were not involved in the design of the study, the collection, analysis and interpretation of data, writing the manuscript or the decision to submit the manuscript for publication.

## Data availability

The subcortical auditory segmentations of the study-specific template (as well as T1-weighted, DEC-TDI, and WM FOD templates) are available on the Open Science Framework: https://osf.io/pxmf5/.

